# Molecular evolution in small steps under prevailing negative selection – A nearly-universal rule of codon substitution

**DOI:** 10.1101/510735

**Authors:** Qingjian Chen, Ao Lan, Xu Shen, Chung-I Wu

## Abstract

The widely accepted view that evolution proceeds in small steps is based on two premises: i) negative selection acts strongly against large differences (Kimura 1983); and ii) positive selection favors small-step changes (Fisher 1930). The two premises are not biologically connected and should be evaluated separately. We now extend the approach of Tang et al. (2004) to codon evolution for the entire genome. Codon substitution rate is a function of the physico-chemical distance between amino acids (AAs), equated with the step size of evolution. This step size depends on a large number of physico-chemical properties as 46 of the 48 properties examined affect the rate. Between 9 pairs of closely-related species of plants, invertebrates and vertebrates, the evolutionary rate is indeed strongly and *negatively* correlated with the AA distance (Δ_U_, scaled to [0, 1]). While the analyses corroborate the published results that relied on partial genomes, there is an important difference: Δ_U_ is strongly correlated with the evolutionary rate (R^2^ > 0.8) only when the genes are under predominant negative selection. Nevertheless, since most genes in most taxa are strongly constrained by negative selection, Δ_U_ would appear to be a nearly-universal measure of codon evolution. In conclusion, the driving force of the small-step evolution at the codon level is negative selection. The unanswered question of whether positive selection may, or may not, follow the small-step rule will be addressed in a companion study (Chen, et al. 2019).

## Introduction

Since the time of Darwin, biologists have accepted that evolution proceeds in small steps. An obvious explanation is mutational input: large changes require many mutations and each only makes an incremental contribution. Approaching the issue from the angle of natural selection, R. A. Fisher formalized the selectionists’ view of small-step evolution, known as Fisher’s Geometric Model (FGM) (Fisher 1930). FGM uses the metaphor of climbing the adaptive peak in a multi-dimensional landscape and suggests that small changes are more likely to be advantageous than large ones. At the molecular level, the neutral theory also posits small-step evolution, which can be summarized by two rules (Kimura 1983). 1) Functionally less important molecules evolve faster than functionally important ones. 2) Variants that are functionally similar to the wildtype are more likely to be substituted than dissimilar ones. The two rules are mainly about escaping negative selection but, when applied to positive selection, would converge with the FGM view. In this study, we focus on the second rule - coding-sequence evolution in small steps.

It is noteworthy that selectionists and neutralists appear to agree on small-step evolution, albeit with different emphases. In the neutralists’ view, negative selection tolerates small-step changes while FGM postulates that positive selection favors small-step improvements. Because negative and positive selection are distinct forces driving different processes (Wang, et al. 2017), small-step evolution can be considered two models in one.

In this study, the step size of evolution is represented by amino acid (AA) differences (or distances) (Grantham 1974; Dayhoff, et al. 1978; Miyata, et al. 1979; Henikoff and Henikoff 1992; Kumar, et al. 2009; Adzhubei, et al. 2013). One approach to AA distances attempts to identify physicochemical properties of AAs that can best explain long-term substitution patterns. Earlier methods by Grantham (Grantham 1974) and Miyata (Miyata, et al. 1979) and the more recent ones including SIFT (Kumar, et al. 2009) and PolyPhen (Adzhubei, et al. 2013) take this approach. The second approach attempts to identify substitution patterns directly from protein or DNA sequences and uses these evolutionary patterns as the proxy for AA distances. This second approach can be either amino acid-based or codon based. For example, PAM (Dayhoff, et al. 1978) and BLOSUM (Henikoff and Henikoff 1992) are AA-based, searching for long-term evolutionary patterns among the 190 (= 20 × 19/2) pairwise comparisons. In contrast, the codon-based approach (Yang, et al. 1998; Tang, et al. 2004; Tang and Wu 2006) compares closely-related species whose triplet codons differ by at most 1 bp. Among the 190 pairs, only 75 pairs can be exchanged by a 1-bp mutation. We shall take the codon-based approach to AA distances (Tang, et al. 2004; Yang 2007).

The study, extending and revising the results of Tang et al. (2004) and Tang and Wu (2006), uses the tools developed therein. The extensions are in three directions. First, in earlier studies, partial genomes are used. Given the partitions of AA changes into 75 kinds, the statistical resolution was barely adequate. These early releases of genomic data are also biased toward functionally important genes that may bear distinct signatures of selection. Hence, whole-genome sequences obtained in the intervening years should be most useful. Second, it is important to densely sample different taxonomic ranks (such as vertebrates, mammals and primates) to test the consistency within the same phylum, class or order. Third, and most important of all, previous studies did not separate the effects of positive and negative selection. For that reason, deviations from the general rule, potentially most informative about the working of the two opposing forces, have been ignored in previous studies.

With the AA distance as the step size of molecular evolution, the effects of negative and positive selection are separately analyzed in this and the companion report (Chen, et al. 2019). Here, we ask whether negative selection drives small-step evolution and whether there exist common rules at the codon level across a wide range of taxa.

## Results

By convention, the number of nonsynonymous changes per nonsynonymous site is designated Ka (Kimura 1983; Li, et al. 1985)(or dN,(Nei and Gojobori 1986)) and the corresponding number for synonymous changes is Ks. Ka is thus the aggregate measure of all nonsynonymous changes and can be decomposed into 75 classes of amino acid (AA) substitutions whose codons differ by only one bp. Each of these classes is labeled Ki, i=1, 75. Hence,

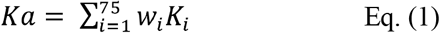

where w_i_ is the weight reflecting the number of sites available for AA exchanges of the i-th pair. In the PAML software package (Yang 2007), Ki’s are obtained first when calculating Ka by Eq. (1). Therefore, no additional calculation is needed. In the next two sections, we will show that i) Ki’s are nearly universally correlated across taxa; and ii) This high correlation reflects the physicochemical properties of AAs.

### I. The correlation among Ki’s across taxa

We first calculate Ki between nine pairs of species using their whole-genome sequences (Fig. 1A). Among them are one pair from plants (*Arabidopsis*), one pair of insects (*Drosophila*), and seven pairs of vertebrates with rodents (*Mus - Rattus*) representing the vertebrates. Each pair consists of two closely related species. Genome-wide Ka/Ks ratios of these nine pairs ranges from 0.12 to 0.27.

**Figure 1.**
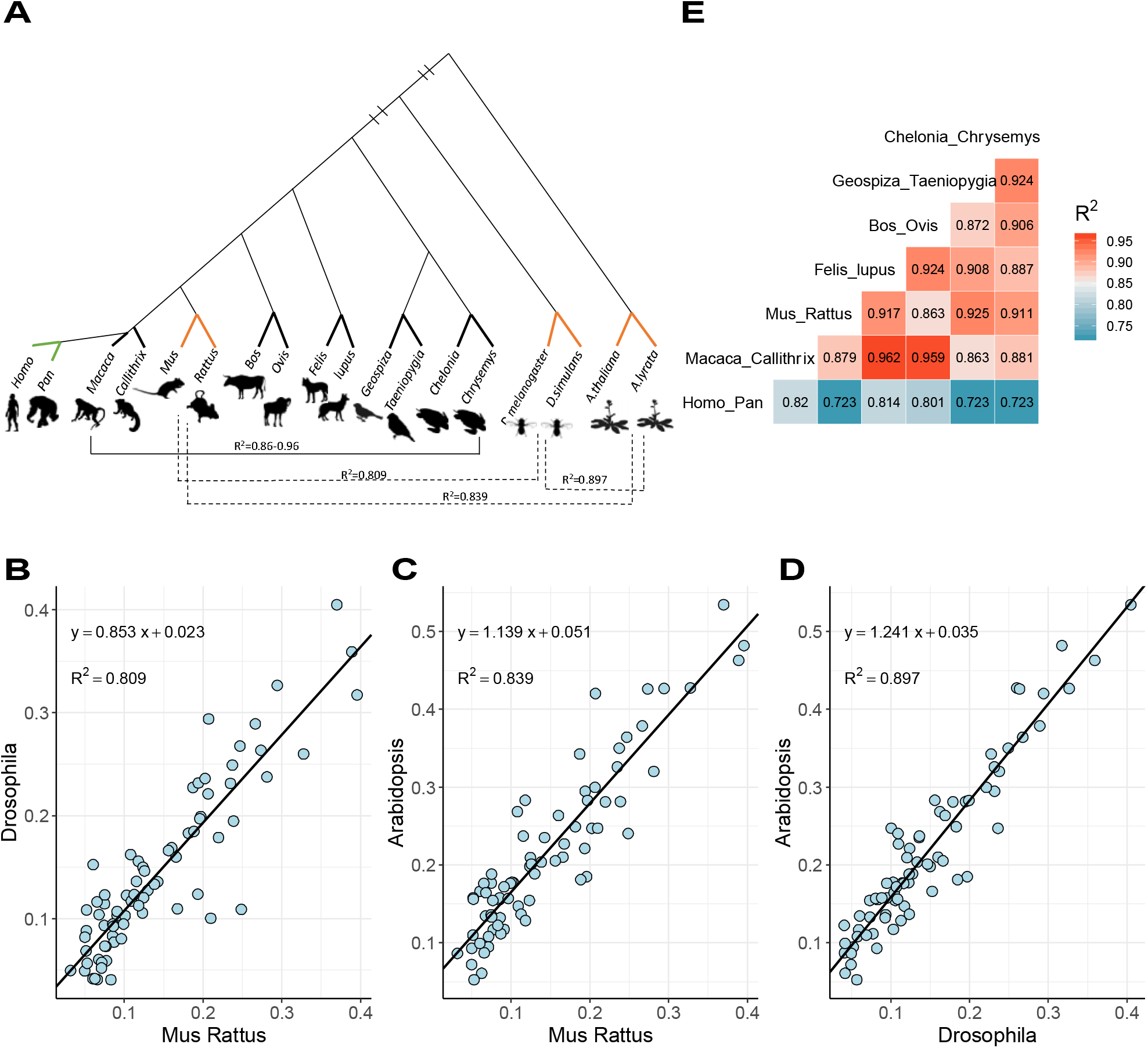
Correlation of Ki/Ks values between different taxa. **(A)** Phylogenetic Tree of 9 representative species pairs (full names are shown in Table S1). R^2^ values (pairwise correlation of Ki’s) range from 0.86 to 0.96 in 6 vertebrates (horizontal solid line), except for Homo-Pan pairs. Three species pairs, Rodent, Drosophila and Arabidopsis (bold orange lines), are selected to show species pairs with long evolutionary distance. **(B-D)** Linear regression results of Ki/Ks between Rodent, Drosophila and Arabidopsis. **(E)** Pairwise R^2^ values of Ki/Ks among 7 species pairs in vertebrates.

We then calculate the correlation in Ki between two pairs of distantly related taxa (Fig. 1A). Pairwise correlations between *Arabidopsis (A. thaliana* vs. *A. lyrata), Drosophila (D. melanogaster* vs. *D. simulans*) and vertebrates (*Mus* vs *Rattus*) are shown in Fig. 1B-D. The R^2^ values are 0.81 (vertebrates vs. insects), 0.84 (vertebrates vs. plants), and 0.90 (insects vs. plants). Such strong correlations over a large phylogenetic span suggest that AA substitutions follow a nearly universal rule. While Ki within each taxon may be very different, their relative magnitudes remain the same across all taxa. The correlations of Fig. 1B-1D are close to the results obtained by Tang et al. (2004). Apparently, the early releases of genic sequences are not seriously biased in their evolutionary rates, thus permitting further analyses based on the earlier results.

We now examine the correlation in a dense phylogenetic framework by comparing species pairs between vertebrate classes and between mammalian orders (Fig. 1E). While one might have expected the correlation to be even greater in the lower taxonomic rank, the observations of Fig. 1E suggest otherwise. In fact, the R values are not strongly dependent on the phylogenetic distance. For example, R^2^ in the *Drosophila - Arabidopsis* comparison is higher than many of the 21 comparisons between vertebrates. In particular, the AA substitutions between hominoids and other vertebrates often yield R^2^ < 0.8. It seems plausible that similar forces work reiteratively from taxa to taxa.

The high correlation among Ki values permits a generalized (or universal) Ki measure as proposed before (Tang, et al. 2004). This universal measure, referred to as Ui (Tang, et al. 2004), ranges between 0.25 and 2.5 for i= 1 to 75. Ui is scaled such that the weighted mean is 1 across the 75 classes. For any species, the expected Ki is

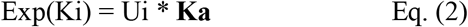

where the boldface Ka denotes the species’ genome-wide value. As long as we know **Ka**, Exp(Ki) can be calculated for i = 1, 75 by Eq. (2).

### II. Ki in relation to AA properties

The high correlation across a large phylogenetic distance (Fig. 1) suggests something as basic as physicochemical properties of amino acids to be a major cause. We thus analyze 48 physicochemical, energetic, and conformational distances among AAs (Fig. 2). We use Ui (i = 1, 75 (Tang, et al. 2004)) to represent Ki across species, as Ui is highly correlated with each species’ Ki (see Fig. 3 below).

**Figure 2.**
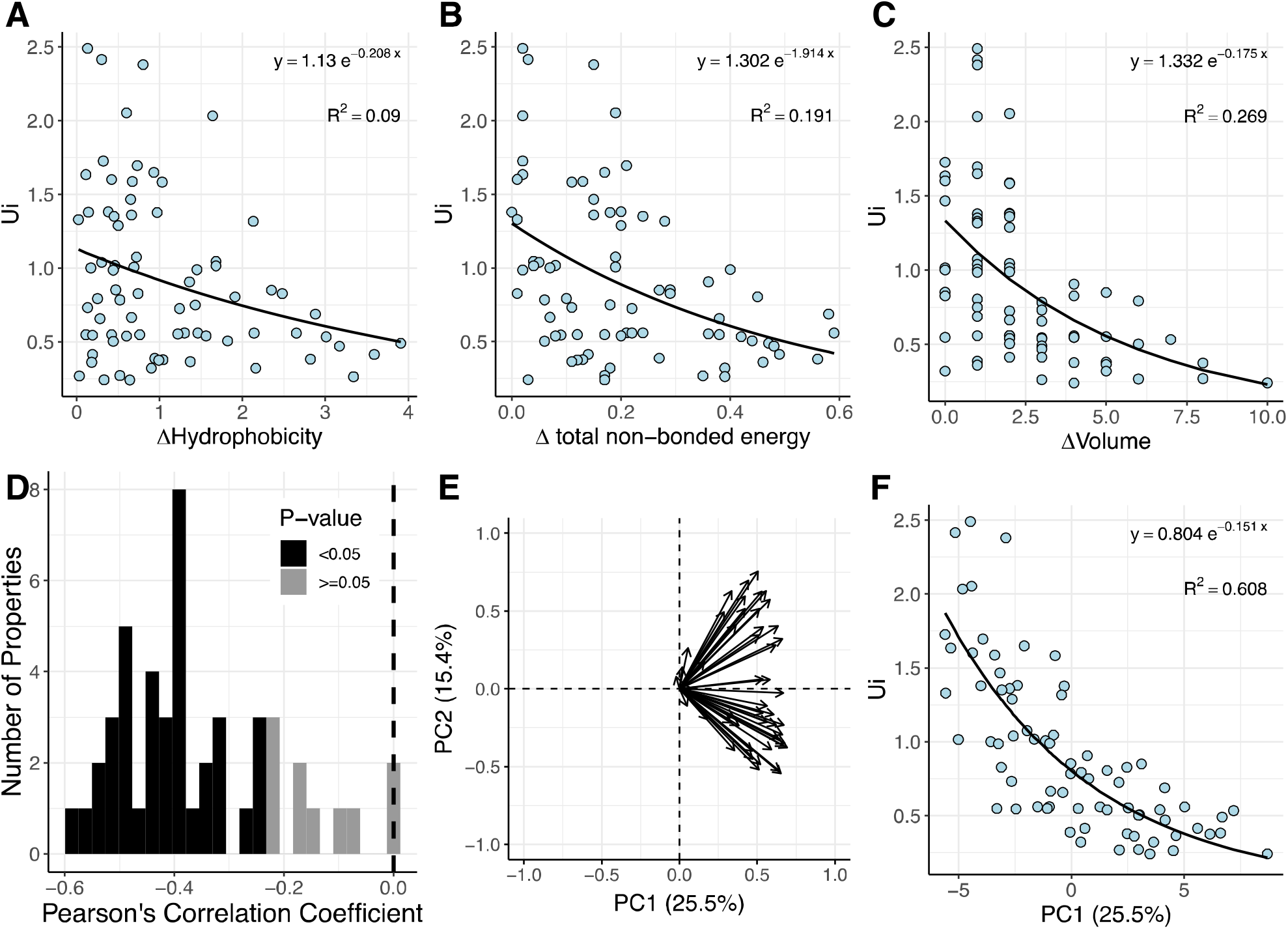
The relationship between physicochemical AA distance and Ui. 48 physicochemical, energy and conformational properties are used in the analysis. **(A-C)** Scatter plot of three selected properties against Ui, including hydrophobicity**(A)**, total non-bounded energy**(B)** and volume**(C). (D)** Pearson’s correlation coefficients between 48 properties and Ui. Among those properties, 46 properties are negatively correlated with Ui, where 38 of them are significantly (black, p-value < 0.05). **(E)** Contribution of 48 properties to the first two principal components. Properties cluster into two categories in the second principal component but are indistinguishable in the first principal component. **(F)** Scatter plot of first principal component (PC1) against Ui.

**Figure 3.**
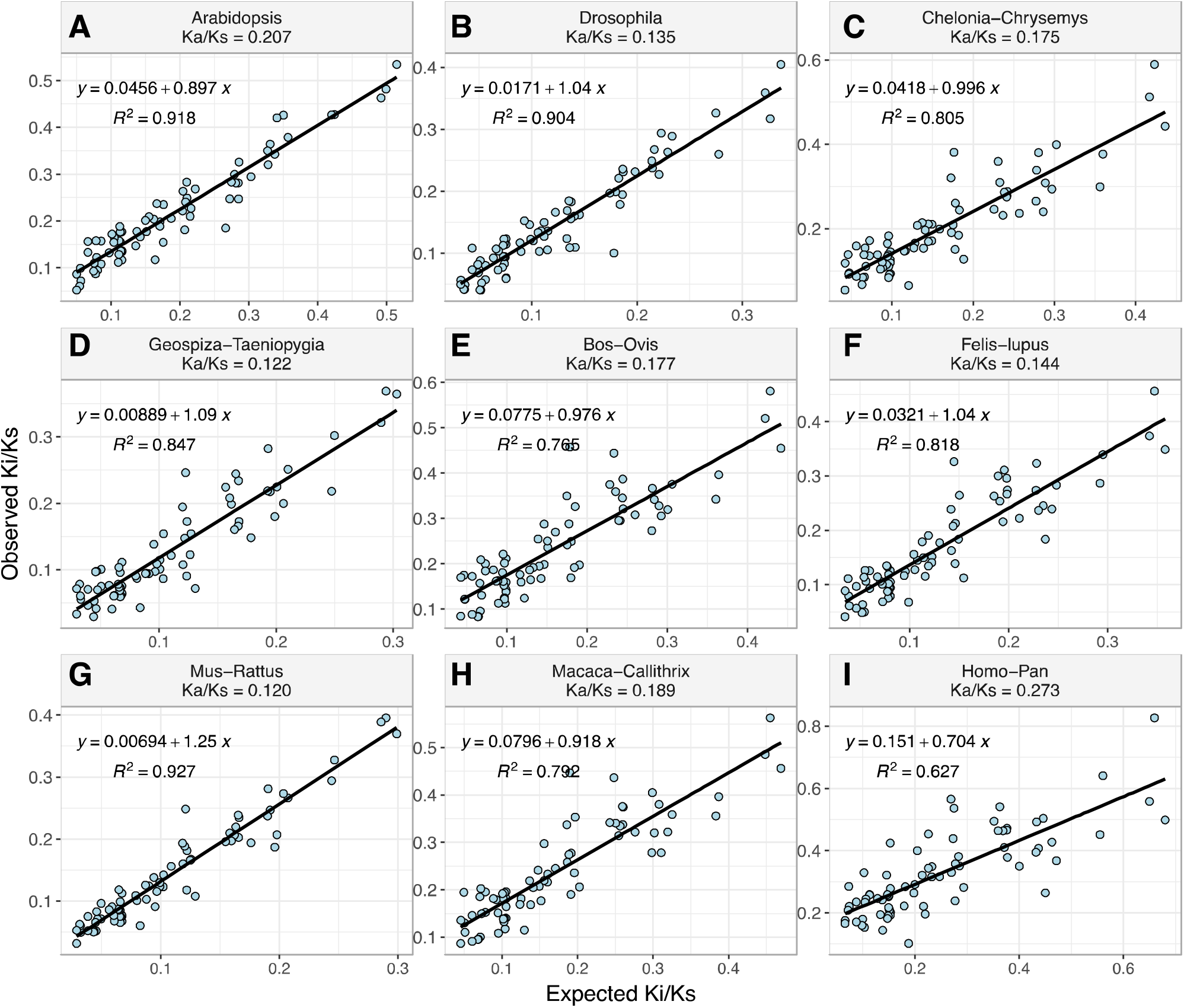
Correlations of expected and observed Ki/Ks for nine species pairs. Since U_i_ = U_1_ - (U_1_ - U75) ΔU_i_, the X-axis label can be written as Exp(Ki/**Ks**) = U_i_\***Ka/Ks** = [2.5 - 2.25 ΔU_i_]\***Ka/Ks**, which is a linear function of ΔU_i_. The boldface **Ka/Ks**, the genome-wide Ka/Ks, is the characteristic of each pair of species.

Ui is fitted to each of the 48 distance measures via an exponential function (see Kimura 1983)(Kimura 1983). Figs. 2A-C show regressions of Ui on hydrophobicity, total non-bounded energy, and volume. Although hydrophobicity is generally thought to be an important functional attribute, the correlation is weak (R^2^ = 0.09). Instead, the volume of AAs explains more of the Ui variation with R^2^ = 0.27. The correlation coefficient for each AA property is given in Fig. 2D, which shows that 46 of the 48 measures are negatively correlated with Ui and P < 0.05 is found for 38 of them (Table S2). Given that only two AA properties are uncorrelated with the evolutionary rate, the fitness effect is quite broadly distributed among many AA properties.

The complexity of translating evolutionary rate and fitness back to biochemistry is further revealed by a Principal Component Analysis (PCA). Fig. 2F shows that PC1 explains about 61% of the Ui variation. Further details can be seen in Fig. 2E. It shows that the contributions to PC1 are distributed broadly among the 48 measures with “helical contact area” in the lead, contributing about 4%.

### III. The general rule for small-step evolution expressed as Δ_U_(i)

Given Ui, the AA distance of the i-th pair can be more conveniently re-scaled as Δ_U_(i) =(UrUi)/(UrU75), which falls in the range of [0, 1] with Δ_U_(1) = 0 for the closest pair, [Ser-Thr], and Δ_U_(75) = 1 for the most distant pair, [Asp-Tyr].

The observed vs. expected Ki for the nine pairs of species are shown in Fig. 3. The X-axis is Exp(Ki/Ks), which can be written as [2.5 - 2.25 ÁU¡] * **Ka/Ks** (see the legend of Fig. 3 and Table S1 for details). There are three groups in this figure. The first group consists of three pairs of well-assembled genomes - *Arabidopsis, Drosophila*, and rodents (Fig. 3A, B, and G). All three taxa show R^2^ > 0.9 between Obs(Ki) and Exp(Ki). It is notable that the R^2^ improves significantly in *Drosophila*, rising from 0.706 to 0.904 in comparison with Tang et al. (2004), as the number of genes increases from 309 to 9,710. In the second group of five pairs of genomes of moderate quality, R^2^ ranges between 0.77 and 0.85. Taking into account the room for improving their quality, we conclude that these species follow the nearly universal pattern of codon substitutions.

The third group consists of the one exceptional case of human-chimpanzee comparison, which yields an R^2^ of 0.627, far lower than the rest. There may be several explanations for this unusually low R^2^. Since codon substitutions involving CpG sites have been removed from consideration (see Methods), this obvious explanation is ruled out (see Fig. S1). The second explanation is the unusual selective pressure in the human lineage (Bustamante, et al. 2005; Williamson, et al. 2005; Subramanian 2011). However, because R^2^ along the human, chimpanzee, and gorilla lineages is, respectively, 0.572, 0.638, and 0.646 (Fig. S2 and Supplementary Note), the human lineage does not stand out in this respect. Instead, we will show that hominoids as a group are unusual and many genes in their genomes have a high Ka/Ks ratio > 0.6.

### IV. The rule of small-step evolution is governed by strong negative selection

Between each pair of species, we divide the genomes into five bins in the ascending order of Ka/Ks (see Methods). The Ka/Ks of each bin is plotted against the R^2^ value [i.e., Obs(Ki) vs. E(Ki)] of that bin in Fig. 4. There are two obvious trends. First, R^2^ is generally > 0.6 for gene groups with Ka/Ks < 0.4. The drop in R^2^ become steeper as Ka/Ks gets larger than 0.4. This important trend appears to escape notice in Tang et al. (2004). Second, the human-chimpanzee comparison follows the same pattern but generally at a lower level.

**Figure 4.**
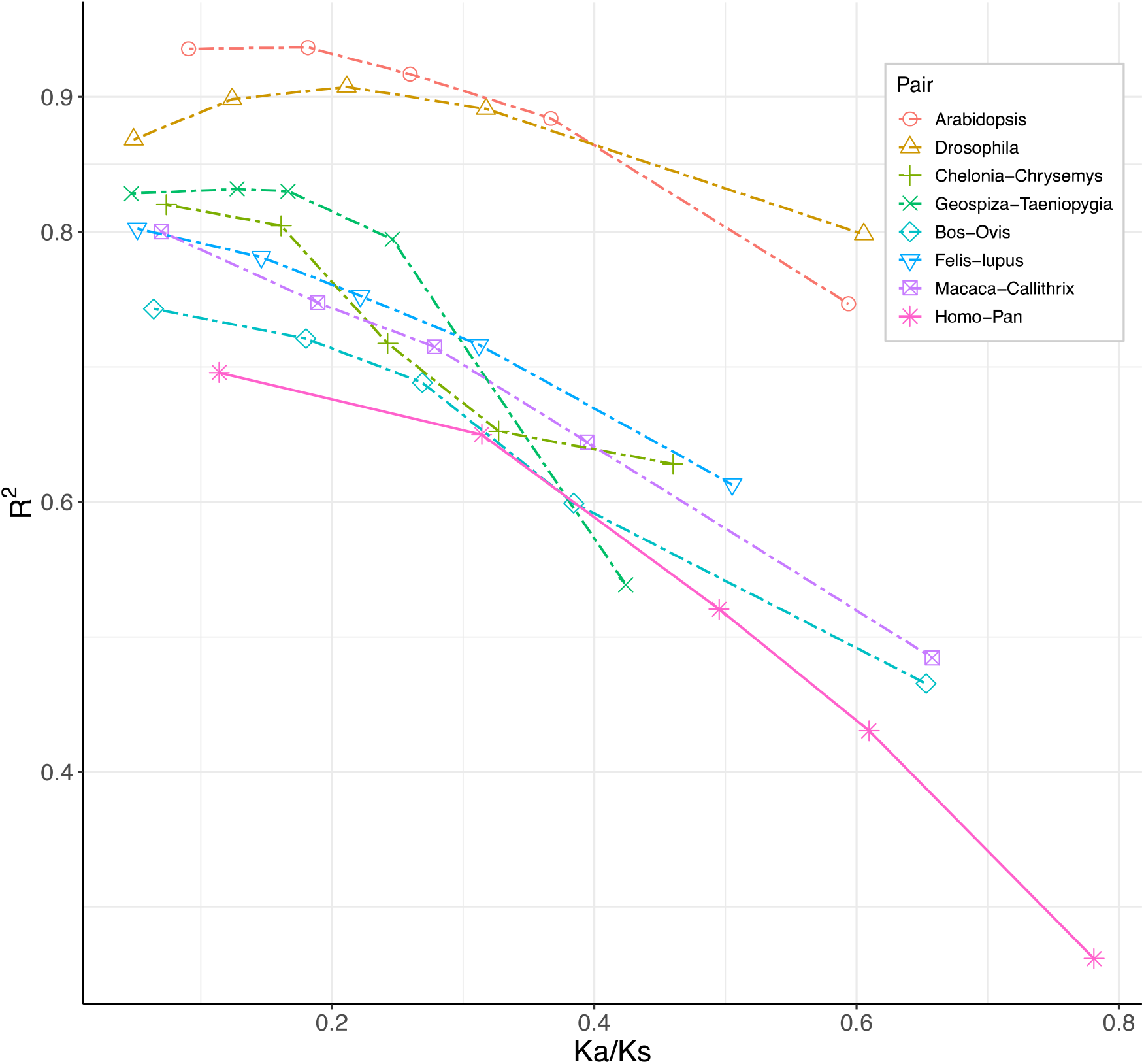
Relationship between R^2^ values(Obs(Ki) vs. Exp(Ki)) and selection strength(Ka/Ks). Orthologous genes in each pairs of species are divided them into 5 categories with equal non-synonymous changes, according to their Ka/Ks ratio. The x-axis is average Ka/Ks ratio of each category in each species pair and the y-axis is the R^2^ value(squared correlation coefficient) of their expected against observed Ki/Ks.

There are two opposing forces driving the trend of Fig. 4 when Ka/Ks increases: weaker negative selection and/or stronger positive selection. Separating the two effects will be the subject of the accompanying study (Chen, et al. 2019). Where Ka/Ks < 0.4, we can nevertheless draw the following conclusion: The evolution of genes under strong negative selection takes small steps and follows a nearly universal rule. This rule is governed by the physicochemical properties of AAs.

## Discussion

The average pattern of molecular evolution is driven predominantly by negative selection. In most genes of most taxa, we obtain Ka/Ks < 0.3, which means the elimination of more than 70% of non-synonymous mutations. The prevailing negative selection has led to a strikingly simple pattern: codon evolution takes small steps and follows a nearly universal rule. By this rule, when an exchange between AA1 and AA2 is five times more likely than that between AA3 and AA4 in mammals, the same ratio would be preserved in non-mammalian vertebrates, invertebrates, and plants. The relative magnitude remains nearly constant.

Given the general pattern in such diverse taxa, negative selection at the codon level must be operating at a basic level of biochemistry. We show that the working of negative selection depends on the properties of the AA residues and almost all measures of AA properties contribute to the fitness difference. Even the most obvious properties like hydrophobicity and non-bonded energy contribute only a small fraction to the overall evolutionary rate. Collectively, the first principal component among 48 properties explains about 60% of the rate variation. There may still be many missing measurements.

The simple evolutionary pattern associated with the complex biochemistry provides an important lesson on the physicochemical basis of traits and diseases. There have been many proposals for measuring AA distance as an index of fitness difference (Grantham 1974; Dayhoff, et al. 1978; Miyata, et al. 1979; Henikoff and Henikoff 1992). Genetic disease associated with AA changes is such an example of negative selection (Adzhubei, et al. 2013). Since matrices relying on a few biochemical properties are not likely to capture much of the evolutionary pattern (see Fig. 2), it may be more informative to assess the evolutionary rate directly from DNA sequence data (Hanada, et al. 2007). In this perspective, the Δ_U_ measure of codon substitution should be particularly suited to investigating the strength of negative selection.

In comparison with previous studies that link AA properties with molecular evolution at the codon level (Zuckerkandl and Pauling 1965; Epstein 1967; Clarke 1970; Grantham 1974; Miyata, et al. 1979; Kimura 1983; Tang, et al. 2004), this study shows clearly the need to separate the effects of negative and positive selection. While the working of negative selection is, to some extent, predictable, positive selection may show very different patterns, which will be addressed in the accompanying study (Chen, et al. 2019). Indeed, some previous studies have reported unusual AA substitution patterns in extreme environments or under domestication, where positive selection could be prevalent (Lu, et al. 2006; Luo, et al. 2017; Xu, et al. 2017).

## Materials and Methods

### Multiple Alignment Data

A multiple alignment file of 99 vertebrates with human for CDS regions was downloaded from the UCSC Genome Brower (http://hgdownload.cse.ucsc.edu/goldenPath/hg38/multiz100way/alignments/knownCanonical.exonNuc.fa.gz). We then selected seven representative pairs of species across vertebrates (Table S1). There were two pairs of species from the Primates order, one was from the family *Hominidae (Homo sapiens* and *Pan troglodytes*) and the other from the family *Cercopithecidae (Macaca fascicularis* and *Callithrix jacchus*). The other five pairs of species were from the order *Rodentia (Mus musculus* and *Rattus norvegicus*), the order *Carnivora (Felis catus* and *Canis lupus familiaris*), the order *Artiodactyla (Bos taurus* and *Ovis aries*), the class *Aves (Geospiza fortis* and *Taeniopygia guttata*), and the class *Reptilía (Chelonia mydas* and *Chrysemyspicta bellif*). Additional technical details used here can also be found in Lin et al. (Lin, et al. 2018) and Wang et al. (Wang, et al. 2018).

A multiple alignment file of 26 insects with *D. melanotaster* was also downloaded from the UCSC Genome Brower(http://hgdownload.cse.ucsc.edu/goldenPath/dm6/multiz27way/alignments/refGene.exonNuc.fa.gz). The genome sequences of *D.melanogaster* and *D.simulans* were then extracted from this file.

To correct for multiple hits for Ki/Ks, two closely-related species of each pair were chosen from the same family. The alignment files were filtered using the following criteria: 1) Filter out genes in un-canonical scaffolds. 2) Use the corresponding amino-acid multiple alignment file to remove non-coding genes. 3) Filter redundant transcripts generated by alternative splicing, based on a gene-transcript id conversion table downloaded from Ensembl biomart (www.ensembl.org/biomart). 4) Genes aligned with too many gaps (more than 5%) were removed. 5) Genes shorter than 90 bp were removed. 6) Mutations related to CpG sites in the Homo-Pan pair were masked because CpG-related mutations account for more than 30% of total mutations in this pair. Hence, we masked CpG related mutations (CG => TG, CG => CA were replaced with NG => NG and CN => CN respectively) using gorilla as the outgroup (Fig. S1, Supplementary Note).

We first obtained protein and DNA sequences of *Arabidopsis thaliana* and *Arabidopsis lyrata* from the Phytozome database (Goodstein, et al. 2012). We then extracted gene alignments using the PAL2NAL program (Suyama, et al. 2006) with the mRNA sequences and the corresponding protein sequences for 1:1 orthologous pairs aligned by MUSCLE (Edgar 2004) using default parameters.

### Ki calculation

We used the sub-program *“codeml”* (codon-based maximum-likelihood) from PAML to calculate each Ki/Ks (Yang 2007). PAML considers codons as units of evolution. The substitution from codon u to codon v is

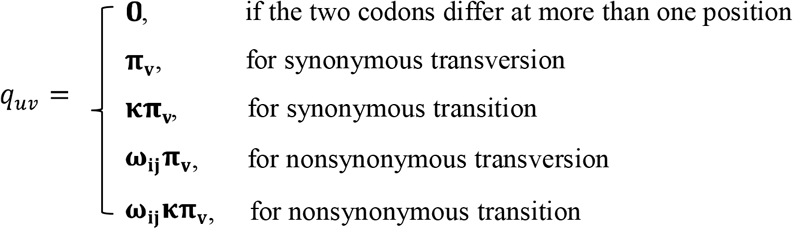

Here, *q_uv_* is the instantaneous rate from codon u to codon v. **k** is the transition/transversion rate ratio, **π**_v_ is the equilibrium frequency of codon v. ω_ij_ is the non-synonymous/synonymous rate ratio, where i (= aa_u_) and j (=aa_v_) are the two amino acids involved. There are 75 ω_ij_, analogous to Ki/Ks (i =1:75), of amino acid pairs whose codons differed by 1 bp. The method considering unequal substitution rate between different amino acids has been summarized in Yang et al. (1998)(Yang, et al. 1998).

The parameters in PAML are as follows: 1) *Model=0*. Use the site model to allow 0) to vary among sites. 2) *codenFreq=0*. Assume individual codon frequencies are equal. 3) *NSsites=0*. Use the M0 (one ratio) model to estimate ω. 4) *aaDist=7*. Allow to estimate amino-acid substitution rates separately. Expected Ki/Ks values were calculated as Exp(Ki/**Ks**) = Ui\***Ka/Ks** in each pair of species.

### Analysis of the physicochemical properties of amino acids (AAs)

Amino acid properties were extracted from Table 2 of Ref. Gromiha et al. (1999) (Gromiha, et al. 1999). In total, 48 selected physicochemical, energetic, and conformational properties are given. The distances between AA pairs were defined by the differences in the raw values shown in their Table 2. Various distance measures yielded similar results. The 48 distances of each AA pair were then scaled by the z-score method for the PCA analysis. PCA was performed using the “factoextra” R package (Kassambara and Mundt 2017). Exponential fitting was accomplished by using the function ‘nls’ in R.

### The relationship between Ui and Ka/Ks

Orthologous genes for each paired species were divided into five categories with approximately equal number of non-synonymous changes according to their Ka/Ks ranking. Genes with fewer than two mutations were removed. In an ideal case, Ka values increase when Ka/Ks values ascend and Ks values are constant in the five categories. This is true in most of cases. However, the top 20% group in Homo-Pan is an outlier, exhibiting a sharp decrease in Ks. The Ks values of the five categories in the Homo-Pan comparison were replaced by genome-wide Ks. The Mus-Rattus comparison was removed in Fig. 4 because Ui values were deduced from the comparison between Mus-Rattus and a pair of yeast (Tang, et al. 2004).

## Acknowledgements

We would like to thank Ziwen He, Haijun Wen, Hao Yang, Qipian Chen and members of Wu Lab for discussions and advices. This work was supported by the National Natural Science Foundation of China [91731301 and 31730046), the 985 Project [33000-18831107], and National Key Basic Research Program of China [2014CB542006].

