## Supplementary for "Molecular evolution in small steps under prevailing negative selection – A nearly-universal rule of codon substitution"

**Supplementary Note**

**A short review for AA's evolutionary rate and physiochemical properties**

A key argument between neutralist and selectionist is the evolutionary rate between conservative and radical changes. On one hands, for Darwinian evolution, there must be enough difference in the functional changes for natural selection operating, thus one would expect more radical changes than conservative changes in the AA's evolution. On the other hands, if selection is strictly or nearly neutral , physicochemical similar ones would occur more frequently than dissimilar ones.

There is a long history to compare the AA's evolutionary rate and physicochemical properties . The first work was done by Zuckerkandl and Pauling (1965) , they found amino acid with similar physicochemical properties were substituted more frequently than dissimilar ones. The latter woks including Epstein(1967), Clarke (1970), Grantham (1974) both support this observation above . Miyata et al (1979) obtained a higher correlation between the evolutionary rate and amino acid distance by considering substitutions only attributed to 1-bp mutation, same as our work. Kimura (1983) extended this method and fit the physicochemical distance and evolutionary rate by an exponential function Y=A*e*^BX^. He used the Miyata chemical distance as the physicochemical distance and the relative frequency of amino acid substitutions from McLachlan (1972) . He found the B is -0.376 and the correlation coefficient is -0.742 (R^2^ is 0.55 ). This strongly negative correlation between the physicochemical properties and AA's evolutionary rate gives powerful supports for the neutral theory.

In summary, our work extends the concept of neutral theory. Our work first support the physicochemical properties account for more than 60% variance of AA's evolutionary rate. Then If the similar ones occur more frequently than dissimilar ones and the AA's physicochemical properties shared by every organism, evolutionary rate of different AA should also be shared in most of organisms. The nearly universal AA's exchangeabilities across a wide phylogenetic distance support such hypothesis. The AA's evolutionary rate thus follows a simple and universal rule. The AA’s evolutionary rate is predictable by “universal exchangeabilities” given in Tang et al. (2004) if only the average Ka/Ks ratio is not large.

**CpG related mutations affect the accuracy of Ki's estimation**

The CpG related mutations account for more than 30% total mutations in germline, elevating mutation rate both in germline and somatic levels. As PAML takes a consent ts/tv (Transition / Transversion) rate in every sites in calculating Ki's , the elevating of CpG related mutations will overestimate the ts/tv ratio, leading to severe bias when estimate the Ki's between Homo-Pan pair. We note that, when masking CpG related mutations, The R^2^ (Obs(Ki) vs. Exp(Ki)) is 0.627 between Homo-Pan pair, significantly higher than without masking CpG mutations, 0.372 (Fig.S1 A-B). Meanwhile, ts/tv decrease from 4.36 to 3.22, when masking CpG related mutations. The Ki from AAs affected by CpG related mutations within codons, decrease sharply when masking CpG mutations, such as His-Arg(HR), Gln-Arg(QR), Met-Thr(MT), Cys-Arg(CR), Arg-Trp(RW) and so on(Fig. S1 C).

**The human-lineage does not contribute the unusually low R^2^ for the Homo-Pan pair.**

Is it the evolution in the human lineage that makes the Homo-Pan comparison distinct? We thus infer the ancestral sequences between human and chimpanzee (labeled A_HC_) using 6 primates as outgroups. The Ki and Ks values in the A_HC_ – human, A_HC_ – chimpanzee and chimpanzee – gorilla branch can then be determined and the R^2^ values are 0.572, 0.638 and 0.646, respectively. (Fig. S2). Therefore, the unusually low R^2^ value appears to be a hominoid phenomenon as the R^2^ is only slightly lower in the human lineage.

**Table S1 - Summary of Datasets**

| **Species1** | **Species2** | **Orthologs** | **Ks** | **Ka/Ks** | ^a^**R^2^** | ^b^***ts/tv*** |
| --- | --- | --- | --- | --- | --- | --- |
| *Arabidopsis thaliana* | *Arabidopsis lyrata* | 19,488 | 0.146 | 0.207 | 0.918 | 1.955 |
| *Drosophila melanogaster* | *Drosophila simulans* | 9,710 | 0.12 | 0.135 | 0.904 | 2.181 |
| *Chelonia mydas* | *Chrysemys picta bellii* | 5,847 | 0.099 | 0.175 | 0.805 | 2.7 |
| *Geospiza fortis* | *Taeniopygia guttata* | 3,676 | 0.128 | 0.122 | 0.847 | 2.392 |
| *Bos taurus* | *Ovis aries* | 10,771 | 0.08 | 0.177 | 0.765 | 4.355 |
| *Felis catus* | *Canis lupus familiaris* | 11,643 | 0.236 | 0.144 | 0.818 | 3.247 |
| *Mus musculus* | *Rattus norvegicus* | 12,854 | 0.205 | 0.12 | 0.927 | 3.504 |
| *Macaca fascicularis* | *Callithrix jacchus* | 13,289 | 0.145 | 0.189 | 0.792 | 3.799 |
| *Homo sapiens* | *Pan troglodytes* | 11,571 | 0.011 | 0.273 | 0.627 | 3.22 |

^a^ R^2^: the squared correlation coefficients between Obs(Ki) and Exp(Ki)

^b^ ts/tv: the transition/transversion rate ratio

Table S2 – Summary of 48 selected physicochemical, energetic and conformational properties of 20 amino acid residues.

| **No** | **Property** | **Details** | **R ^a^** |
| --- | --- | --- | --- |
| **1** | *R*_f_ | chromatographic index | -0.58 |
| **2** | *N*_s_ | average number of surrounding residues | -0.564 |
| **3** | ΔASA | solvent accessible surface area for unfolding protein | -0.544 |
| **4** | −*T*Δ*S*_c_ | unfolding entropy changes of chain | -0.539 |
| **5** | −*T*Δ*S*_h_ | unfolding entropy change of hydration | -0.525 |
| **6** | v | volume (number of non-hydrogen side chain atoms) | -0.511 |
| **7** | V^0^ | partial-specific volume | -0.51 |
| **8** | Δ*C*_ph_ | unfolding hydration heat capacity change | -0.499 |
| **9** | *N*_l_ | average long-range contacts | -0.495 |
| **10** | *M*_w_ | molecular weight | -0.48 |
| **11** | *C*_a_ | helical contact area | -0.48 |
| **12** | *H*_p_ | surrounding hydrophobicity | -0.479 |
| **13** | *E*_l_ | long-range non-bonded energy | -0.473 |
| **14** | Δ*H*_c_ | unfolding enthalpy changes of chain | -0.441 |
| **15** | Ht | thermodynamic transfer hydrophobicity | -0.437 |
| **16** | *B*_r_ | buriedness | -0.433 |
| **17** | *B*_l_ | bulkiness | -0.431 |
| **18** | ASA_D_ | solvent accessible surface area for denatured protein | -0.428 |
| **19** | *E*_t_ | total non-bonded energy (*E*_sm_+*E*_l_) | -0.427 |
| **20** | *H*_gm_ | combined surrounding hydrophobicity (globular and membrane) | -0.427 |
| **21** | *E*_sm_ | short and medium range non-bonded energy | -0.401 |
| **22** | μ | refractive index | -0.398 |
| **23** | Δ*G* | unfolding Gibbs free energy change | -0.392 |
| **24** | Δ*G*_c_ | unfolding Gibbs free energy changes of chain | -0.391 |
| **25** | *P*_β_ | β-structure tendency | -0.39 |
| **26** | Δ*H* | unfolding enthalpy change | -0.384 |
| **27** | F | mean rms fluctuational displacement | -0.382 |
| **28** | Ra | solvent accessible reduction ratio | -0.381 |
| **29** | −*T*Δ*S* | unfolding entropy change | -0.365 |
| **30** | Δ*H*_h_ | unfolding enthalpy change of hydration | -0.348 |
| **31** | ASA_N_ | solvent accessible surface area for native protein | -0.346 |
| **32** | Δ*G*_h_ | Gibbs free energy change of hydration for protein | -0.312 |
| **33** | *G*_hD_ | Gibbs free energy change of hydration for denatured protein | -0.311 |
| **34** | *H*_nc_ | normalized consensus hydrophobicity | -0.307 |
| **35** | p*H*_i_ | isoelectric point | -0.269 |
| **36** | P | polarity | -0.253 |
| **37** | *G*_hN_ | Gibbs free energy change of hydration for native protein | -0.251 |
| **38** | *P*_c_ | coil tendency | -0.233 |
| **39** | f | flexibility (number of side-chain dihedral angles) | -0.226 |
| **40** | s | shape (position of branch point in a side-chain) | -0.225 |
| **41** | α*c* | power to be at the C-terminal of α-helix | -0.216 |
| **42** | *K*^0^ | compressibility | -0.176 |
| **43** | *P*_t_ | turn tendency | -0.163 |
| **44** | p*K*′ | equilibrium constant with reference to the ionization property of COOH group | -0.15 |
| **45** | α*m* | power to be at the middle of α-helix | -0.093 |
| **46** | α*n* | power to be at the N-terminal of α-helix | -0.067 |
| **47** | *N*_m_ | average medium-range contacts | 0.005 |
| **48** | *P*_α_ | α-helical tendency | 0.007 |

The 48 amino acid properties are extracted from Table 2 of Gromiha et al. (1998).

^a^ Pearson’s correlation coefficients between Ui and amino acid distance estimated by each property.

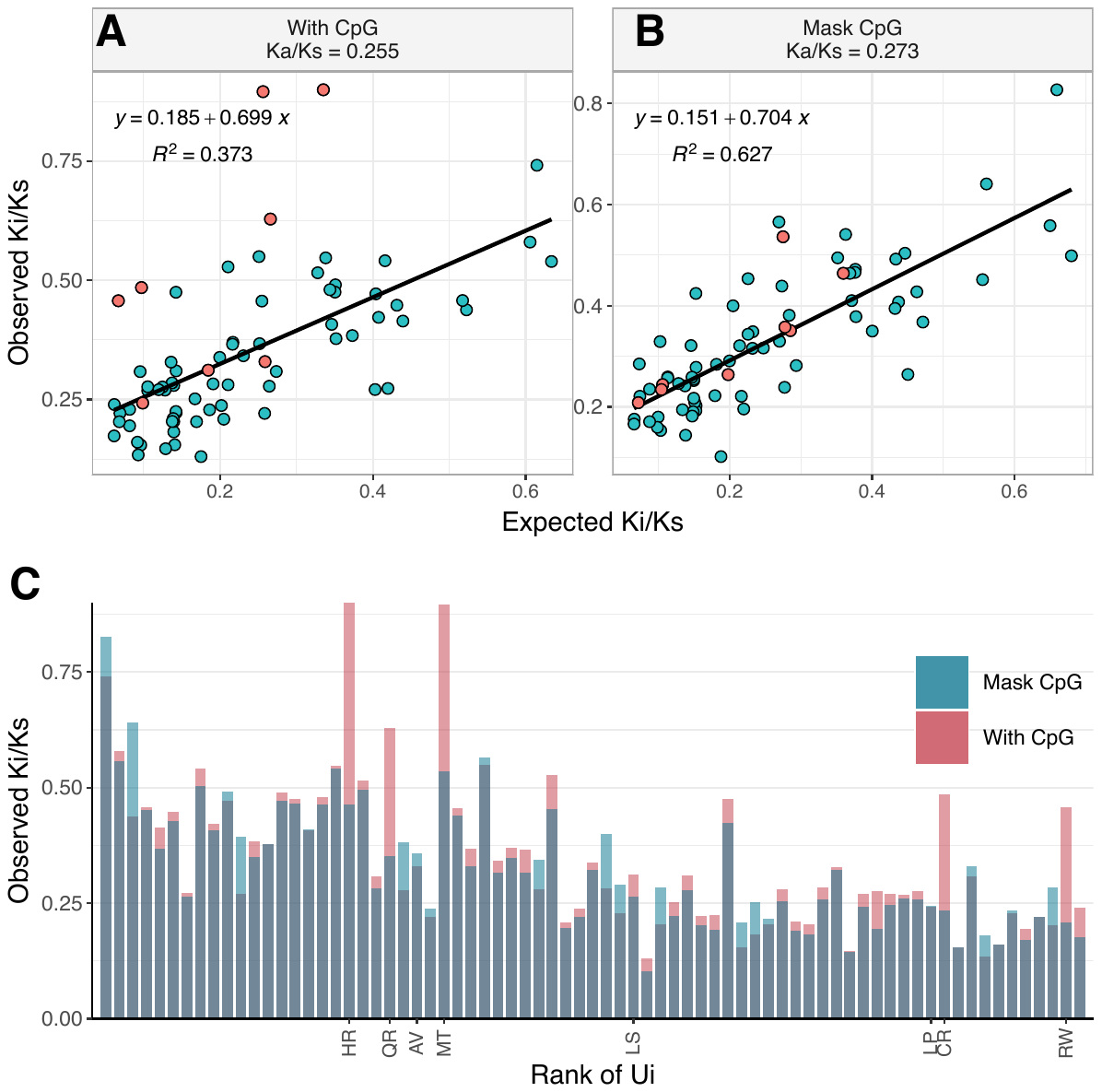

**Figure S1 | Ki/Ks in Homo-Pan pair with and without CpG related mutations.** (**A-B**) R^2^ (Obs(Ki) vs. Exp(Ki)) improves significantly after CpG masking, ascending sharply after masking, from 0.372 to 0.627. Red points are the AAs affected by CpG related mutations within codons (**C**)The Ki/Ks from AAs affected by CpG related mutations within codons, labeled with their one-character abbreviation, decrease a lot after masking CpG related mutations.

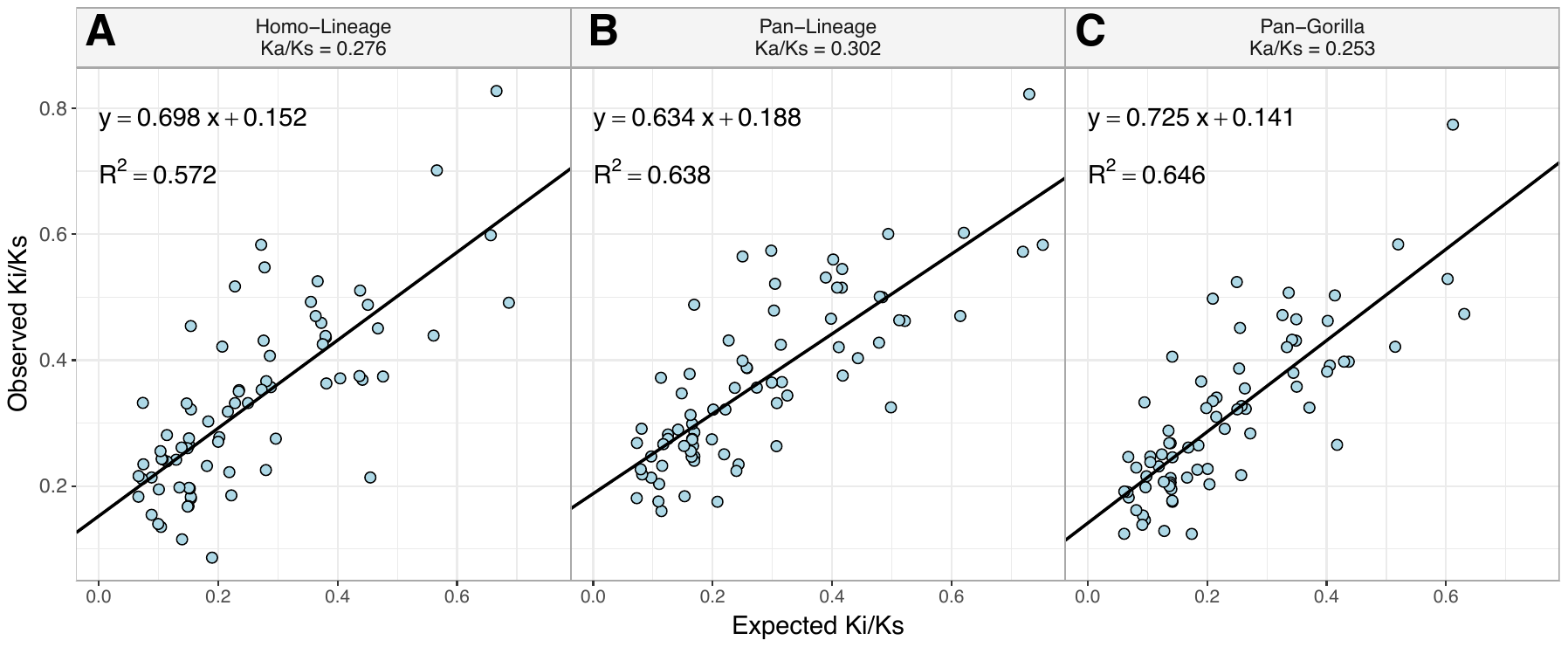

**Figure S2 | Correlations between expected and observed Ki/Ks among apes.** (**A-B**) Homo-Lineage specific and Pan-Lineage specific Ki/Ks was obtained by comparing Homo or Pan sequences with their ancestors.(**C**) The Ki/Ks of pan-Gorilla was obtained by standard Ki/Ks calculation process.
